# Mitomycin C eliminates cyanobacterial transcription without detectable lysogen induction in a *Microcystis*-dominated bloom in Lake Erie

**DOI:** 10.1101/2024.11.06.622312

**Authors:** Robbie M. Martin, Elizabeth R. Denison, Helena L. Pound, Ellen A. Barnes, Justin D. Chaffin, Steven W. Wilhelm

**Author notes:** Address correspondence to Steven W. Wilhelm.

## Abstract

Although evidence indicates that viruses are important in the ecology of *Microcystis* spp., many questions remain. For example, how does *Microcystis* exist at high, bloom-associated cell concentrations in the presence of viruses that infect it? The phenomenon of lysogeny and associated homoimmunity offer possible explanations to this question. Virtually nothing is known about lysogeny in *Microcystis*, but a metatranscriptomic study suggests that widespread, transient lysogeny is active during blooms. These observations lead us to posit that lysogeny is important in modulating *Microcystis* blooms. Using a classic mitomycin C-based induction study, we tested for lysogeny in a *Microcystis*-dominated community in Lake Erie in 2019. Treated communities were incubated with 1 mg L^-1^ mitomycin C for 48 h alongside unamended controls. We compared direct counts of virus-like-particles (VLPs) and examined community transcription for active infection by cyanophage. Mitomycin C treatment did not increase VLP count. Mitomycin C effectively eliminated transcription in the cyanobacterial community, while we detected no evidence of induction. Metatranscriptomic analysis demonstrated that the standard protocol of 1 mg L^-1^ was highly-toxic to the cyanobacterial population, which likely inhibited induction of any prophage present. Follow-up lab studies indicated that 0.1 mg L^-1^ may be more appropriate for use in freshwater cyanobacterial studies. These findings will guide future efforts to detect lysogeny in *Microcystis* blooms.

**Importance:** Harmful algal blooms dominated by *Microcystis* spp. occur throughout the world’s freshwater ecosystems leading to detrimental effects on ecosystem services that are well documented. After decades of research, the scientific community continues to struggle to understand the ecology of *Microcystis* blooms. The phenomenon of lysogeny offers an attractive, potential explanation to several ecological questions surrounding blooms. However, almost nothing is known about lysogeny in *Microcystis*. We attempted to investigate lysogeny in a *Microcystis* bloom in Lake Erie and found that the standard protocols used to study lysogeny in aquatic communities are inappropriate for use in *Microcystis* studies, and perhaps freshwater cyanobacterial studies more broadly. This work can be used to design better methods to study the viral ecology of *Microcystis* blooms.

## Introduction

Harmful algal blooms (HABs) dominated by *Microcystis* spp. occur throughout the world’s fresh waters (1). The detrimental effects of HABs on ecosystems and the services they provide are extensively reviewed (2-5). These detrimental effects have compelled decades of research aimed at understanding the eco-physiology of *Microcystis*-dominated blooms (6, 7).

It is well-accepted that in marine planktonic communities, viruses have significant effects on host abundance, diversity, and distribution (8-10), on host physiology and metabolism (11, 12), food-web function, and biogeochemical cycles (13, 14). A somewhat lesser body of evidence from freshwater studies suggests that viruses have a similar effect on freshwater plankton and are important in the ecology of *Microcystis* blooms (15, 16).

Isolated cyanophages that infect *M. aeruginosa* (17-20) exhibit interactions similar to those seen in marine isolates. For example, viral acquisition of auxiliary metabolic genes of host-like origin indicates that viruses facilitate genetic exchange and influence *Microcystis* metabolism (21, 22). Cyanophage abundance and activity have been correlated to *Microcystis* cell dynamics in blooms (23-25). Metatranscriptomic investigations have further linked viral infection to releases of microcystin into the water column (26, 27). In one case, virus activity is thought to be associated with the 2014 drinking water crisis in Toledo, Ohio (28). More ecologically interesting is that cyanophage infection of *Microcystis* is at least sometimes associated with strain succession over the course of a bloom. An early study linked dynamics of Ma-LMM01-like cyanophages to shifts between toxic and non-toxic strains of *Microcystis* (29). Recent metatranscriptomic studies have confirmed this dynamic while providing deeper community-wide insight (30, 31).

Though evidence suggests cyanophages infecting *Microcystis* are important, questions remain. One question is whether viral activity is the driver or follower of frequently observed replacements of one strain of *Microcystis* with another. Other observations lead to fundamental ecological questions that an examination of *Microcystis* blooms may address. In blooms, *Microcystis* cells can reach concentrations of ∼2 x 10^7^ mL^-1^ (24) and contribute more than 90% of the *in situ* total chlorophyll fluorescence (32). In these circumstances, *Microcystis* seems to violate the precepts of the “Paradox of the Plankton” (33) and outcompetes other taxa of phytoplankton to their near exclusion (34). More confoundingly, *Microcystis* does so in the presence of cyanophages that presumably infect them (30, 31, 35). This latter observation, on its surface, seems to run counter to basic tenets of the “Kill-the-Winner” hypothesis (36). A basic question then becomes, how does *Microcystis* exist at high concentrations in the presence of viruses that infect it?

A possible explanation to this question is lysogeny. Lysogeny is a relationship between a host and a phage where the genome of the virus integrates into the host chromosome (15, 37). The viral genome is then known as a prophage, and the combined virus-cell unit as a lysogen. Prophages can alter gene expression and metabolism of the lysogen through a process termed lysogenic conversion (38). Conversion can enhance lysogen fitness through several mechanisms (39). Applicable to the question at hand is that prophages can offer resistance to infection by closely related phages, a phenomenon termed homoimmunity (40). Thus, lysogen-derived homoimmunity could be an explanation for how *Microcystis* co-exists at high cell concentrations in the presence of cyanophages that can infect it (41).

Little is known about lysogeny in *Microcystis*. Lysogeny in aquatic communities is most commonly tested through chemical induction using the mutagen mitomycin C (42). There is a long history of testing lysogens in marine and freshwater communities using mitomycin C (see Supplemental Table 1 in Knowles et al. (43) for a summary). We found only two studies that attempted induction in *Microcystis* communities. Sulcius et al. (44) incubated colonies collected from a bloom in the Curonian Lagoon in Lithuania in mitomycin C and found no evidence of prophage induction. In an Australian study, bloom communities containing *M. aeruginosa* were treated with mitomycin C, but no evidence of induction was detected (45). However, the same study reported induction of ∼3% of cells in a lab culture of *M. aeruginosa* isolated from a waterbody in Queensland.

While evidence of widespread lysogeny in *Microcystis* during blooms is lacking, a metatranscriptomics study suggested its possibility. Over a period of 5 months in China’s Taihu, Stough et al. (41) observed seasonal patterns in expression of cyanophage genes similar to those of Ma-LMM01 (20). Lytic-associated gene transcripts dominated early in the bloom (June and July), indicating active on-going lytic infections. Dominate expression shifted to lysogenic-associated genes in August through October. Such dynamic shifts in active infections by Ma-LMM01-like cyanophages leads us to posit that lysogeny plays a role in modulating *Microcystis* blooms.

The initial objective of this study was to test for lysogeny in a *Microcystis* bloom. We conducted a microcosm study using natural communities collected in Lake Erie during a *Microcystis*-dominated bloom in 2019. Communities were incubated with 1 mg L^-1^ mitomycin C for 48 h alongside unamended controls. To detect lysogen induction, we compared direct counts of virus-like-particles (VLPs) and examined community transcription for active infection by cyanophage. Mitomycin C eliminated transcription in the cyanobacterial community, while we detected no evidence of prophage induction. Mitomycin C shifted community transcription towards *Alpha-* and *Beta-Proteobacteria*, and subsequent phage infections shifted with these community changes. Inhibition of the target community indicates that use of mitomycin C at 1 mg L^-1^ (the standard protocol) is inappropriate for lysogeny studies in freshwater cyanobacterial communities, and may skew quantitative assessment of other populations.

## Results

### Mitomycin C did not increase VLP abundance in Lake Erie microcosms

In the phosphorus (P)-limited-community experiment, mean VLP concentration was 3.8 x 10^7^ mL^-1^ (SE = 5.3 x 10^6^ mL^-1^) at T_0_. After incubation, there was no difference between groups (ANOVA *p* = 0.78; Figure 1A). In the P-replete-community experiment, mean VLP concentration was 8.2 x 10^7^ mL^-1^ (SE = 1.2 x 10^7^ mL^-1^) at T_0_. After incubation, VLP concentration was lower in the mitomycin C treatment (6.9 x 10^7^ mL^-1^; SE = 2.2 x 10^6^ ml^-1^) than the control (1.5 x 10^8^ mL^-1^; SE = 2.8 x 10^7^ mL^-1^; Tukey’s *p* = 0.04; Figure 1B).

**Figure 1.**
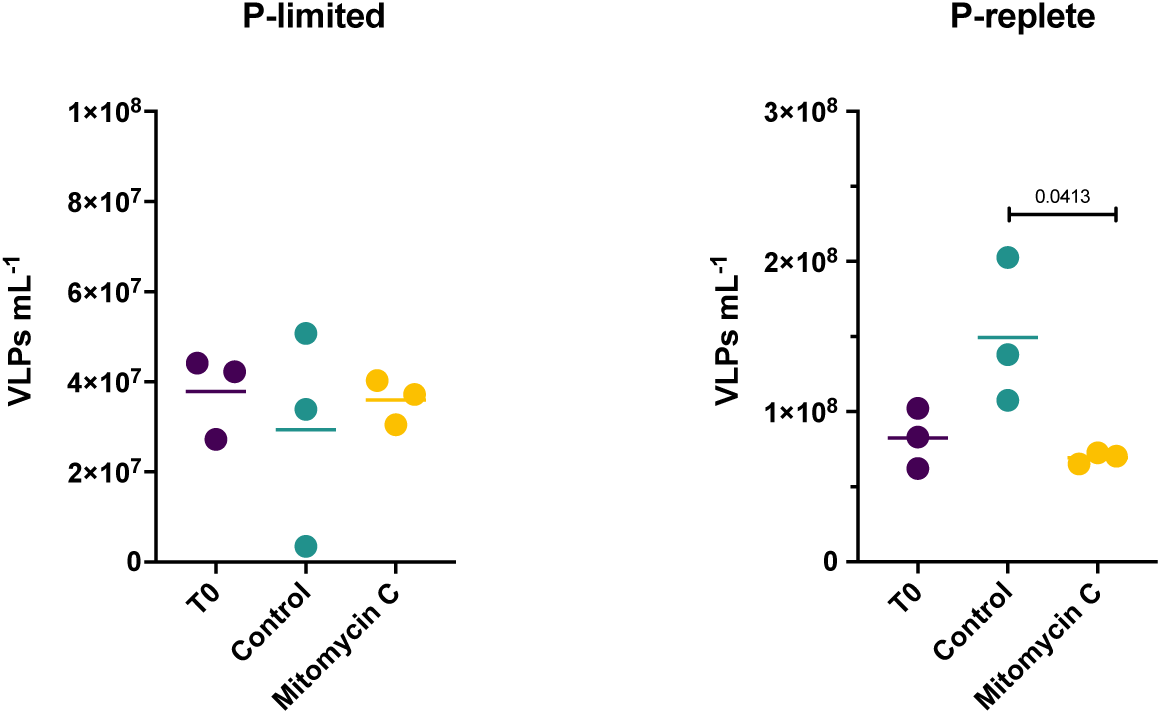
Direct virus counts by treatment. a) P-limited experiment. b) P-replete experiment.

### Sequencing Results

An average of ∼57 M (range 49-68 M) QC reads per library remained for the P-limited experiment, while ∼49 M (range 21-63 M) remained for the P-replete experiment. The co-assembly originating from the P-limited experiment contained 913,786 contigs, within which 1,066,812 putative genes were identified by MetaGeneMark. Of these, 1,061,804 genes were assigned taxonomy by GhostKOALA. The co-assembly of the P-replete experiment contained 553,450 contigs, with 678,427 putative genes identified by MetaGeneMark. Of these, 675,419 genes were assigned taxonomy by GhostKOALA. These taxonomy-assigned gene lists were used for downstream analysis of whole community expression for each of the experiments. For community expression estimation, an average of ∼35 M (61% of QC) and ∼32 M (66% of QC) reads per library mapped to genes of the P-limited and P-replete co-assemblies, respectively.

The VirSorter2/CheckV workflow identified 133 virus contigs in the P-limited co-assembly and 95 in the P-replete co-assembly. In estimating viral expression, an average of ∼225 thousand (0.39% of QC) and ∼172 thousand (0.35% of QC) reads per library mapped to the virus contig lists from the P-limited and P-replete co-assemblies, respectively. Detailed read mapping statistics for both experiments are summarized in Supplemental Table 1.

### Mitomycin C reduced transcriptional representation of Bacteria

Community transcriptional profiles between the two experiments showed similar trends. Mitomycin C only moderately decreased transcriptional representation of Bacteria. This was balanced by increased representation of Plants (primarily green algae) in the P-limited microcosms, and by increased representation across varied taxa in the P-replete microcosms (Figure 2). At T_0_ in the P-limited experiment, Bacteria represented ∼87% of total community expression (Figure 2A). In the control, Bacteria made up ∼81% of total community expression, indicating limited bottle effects. Bacteria representation declined to ∼59% in the mitomycin C treatment (vs. control, Tukey’s *p* = 0.01). At T_0_ in the P-replete experiment, Bacteria represented ∼85% of total community expression (Figure 2B). In the control, Bacteria represented ∼77% of total community expression. Bacteria representation declined to ∼68% in the mitomycin C treatment (vs. control, Tukey’s *p* = 0.08). In the P-limited experiment, chlorophyll *a* (chl *a*) concentration decreased in the mitomycin C treatment vs. control, but not significantly (Supplemental Figure 1A). In the P-replete experiment, chl *a* in the mitomycin C treatment decreased to about half that of the control (*p* = 0.05) (Supplemental Figure 1B).

**Figure 2.**
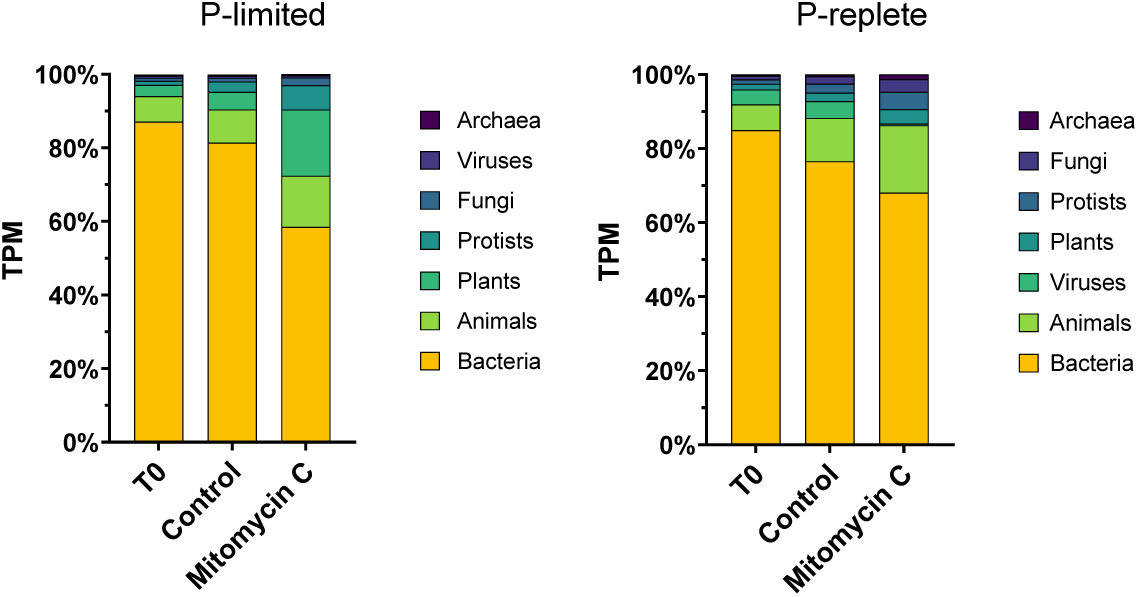
Transcription activity by Kingdom and by treatment as a percent of total community transcription activity. a) P-limited experiment. b) P-replete experiment.

### Mitomycin C effectively eliminated transcriptional representation of Cyanobacteria

Mitomycin C dramatically reduced transcriptional representation of Cyanobacteria and strongly shifted the transcriptional profile of the community toward Proteobacteria and phototrophic Eukaryotes (Figures 3 and 4). In the P-limited controls, Cyanobacteria made up ∼63% of total community transcription, with *Microcystis* alone contributing ∼51% (Figure 3A). In the mitomycin C treatment, Cyanobacterial transcription declined to less than 2% of the community total while *Microcystis* declined to 0.6%. *Microcystis* transcription seemed more heavily suppressed than other Cyanobacteria as it suffered a ∼92-fold decline in transcriptional activity while collectively all other Cyanobacteria declined ∼9-fold.

**Figure 3.**
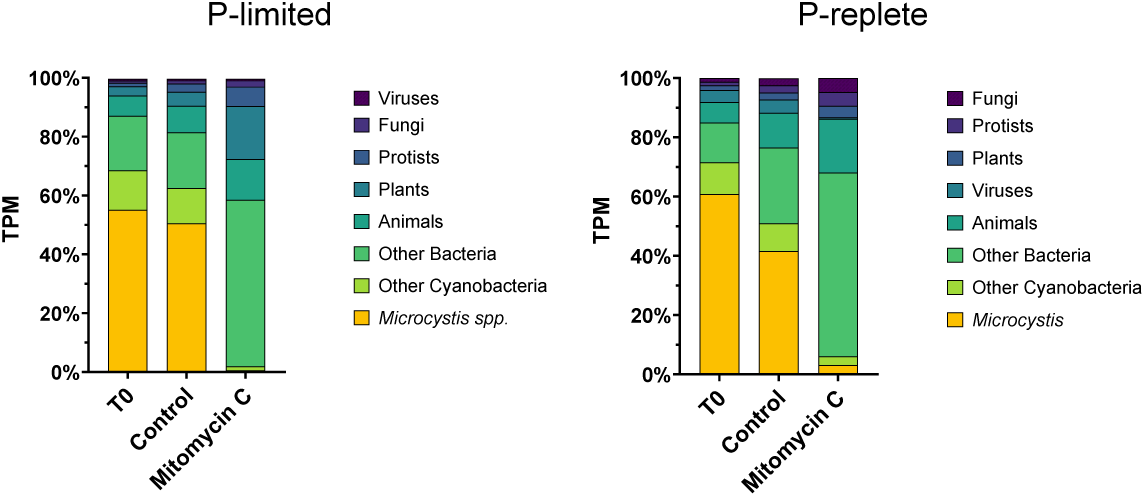
*Microcystis*. transcription activity by treatment as a percent of total community transcription activity. a) P-limited experiment. b) P-replete experiment.

**Figure 4.**
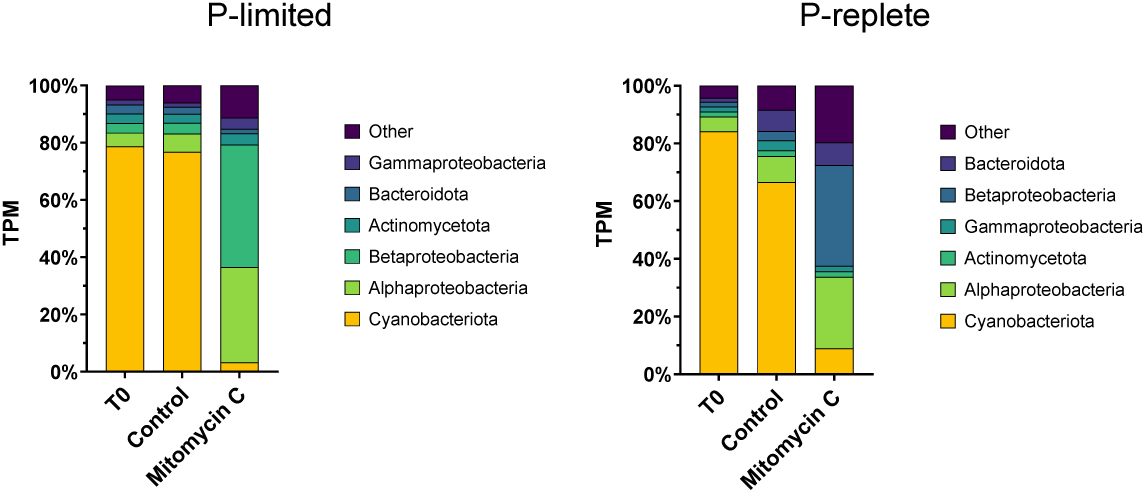
Transcription activity by Phylum or Class of Bacteria and by treatment as a percent of total bacterial transcription. a) P-limited experiment. b) P-replete experiment.

Within the bacterial community proper, Cyanobacteria comprised ∼77% of total Bacteria transcription in the P-limited control and ∼3% in the mitomycin C treatment (Figure 4A). In contrast, transcriptional representation of *Alpha-* and *Beta-Proteobacteria* increased from ∼6% and 4% of the Bacteria total in the control to ∼33% and 43% in the mitomycin C treatment, respectively (Figure 4A).

In the P-replete controls, Cyanobacteria contributed ∼51% of total community transcription, with *Microcystis* alone contributing ∼42% (Figure 3B). In the mitomycin C treatment, Cyanobacteria transcription declined to ∼6% of the community total while *Microcystis* declined to ∼3%. This represents a ∼13-fold decline in *Microcystis* transcriptional activity and only a ∼3-fold reduction among all other Cyanobacteria collectively.

Within the bacterial community proper, Cyanobacteria comprised ∼67% of total Bacteria transcription in the P-replete control and ∼9% in the mitomycin C treatment (Figure 4B). In contrast, transcriptional representation of *Alpha-* and *Beta-Proteobacteria* increased from ∼9% and 3% of the Bacteria total in the control to ∼25% and 35% in the mitomycin C treatment, respectively (Figure 4B). The effects of mitomycin C on major genera of Cyanobacteria, those making >1% of total cyanobacterial transcription in T_0_, is illustrated in Supplemental Figure 2.

### Viral expression followed host changes induced by mitomycin C

Treatment induced clear patterns in viral expression in both the P-limited (Figure 5) and P-replete (Figure 6) microcosms as illustrated in expression heatmaps. Unsurprisingly, expression of major classifications of viruses followed changes in microbial taxa serving as potential hosts.

**Figure 5.**
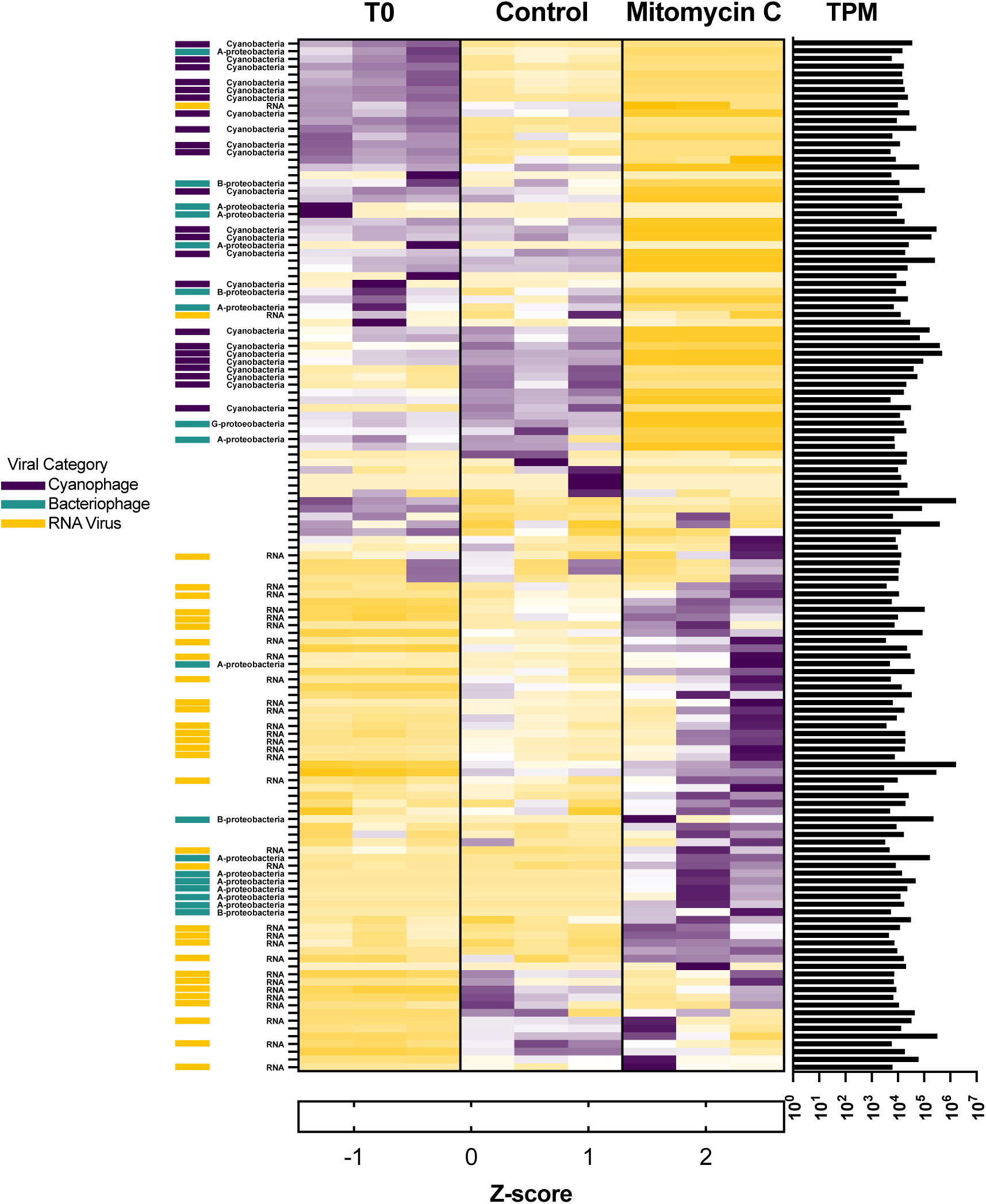
Standardized viral expression in P-limited microcosms. Each row represents expression (TPM) standardized across treatments. Standardized expression for each biological replicate is shown for each treatment.

**Figure 6.**
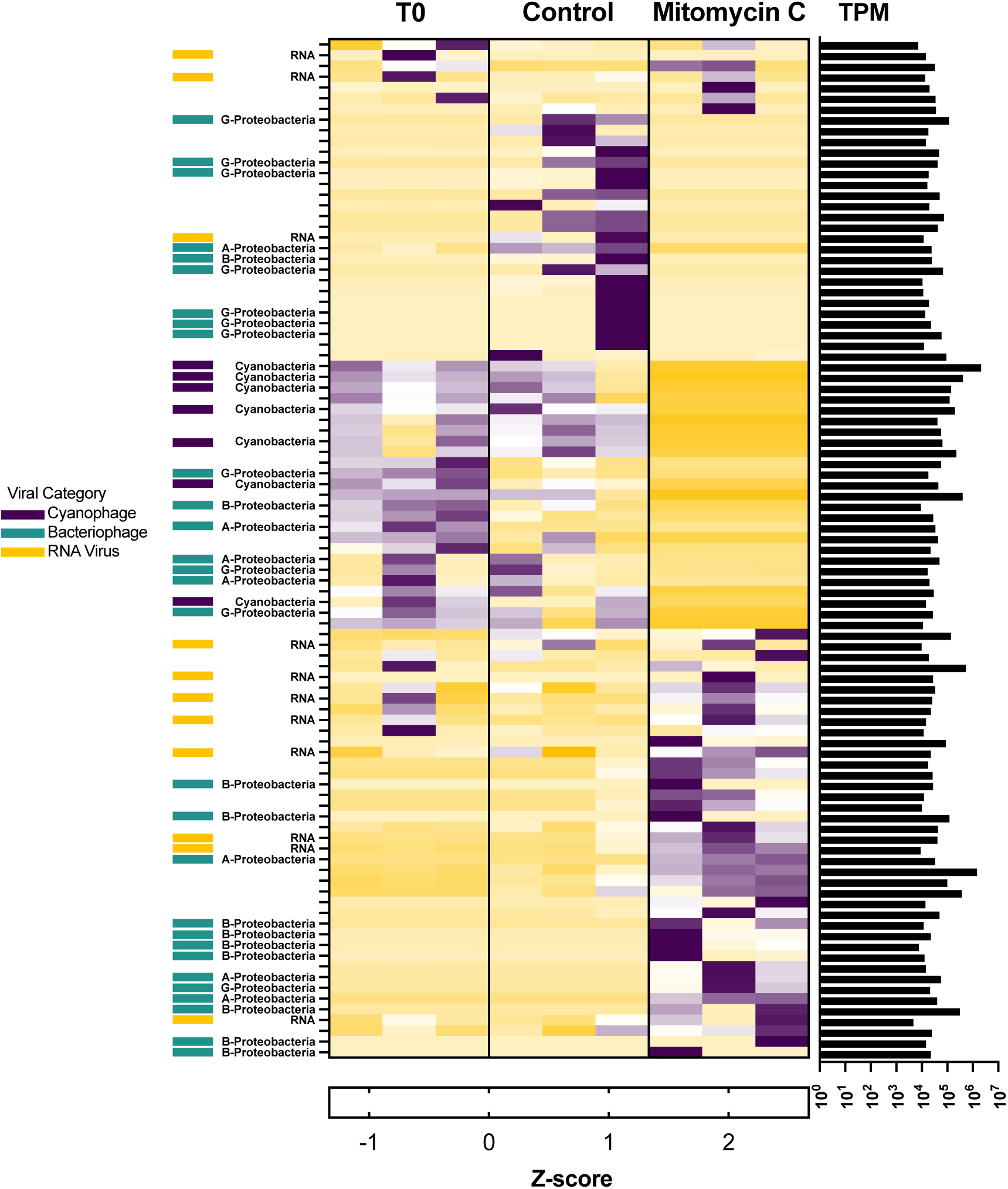
Standardized viral expression in P-replete microcosms. Each row represents expression (TPM) standardized across treatments. Standardized expression for each biological replicate is shown for each treatment.

In the P-limited microcosms, viral expression in T_0_ bottles was dominated by viruses of the phylum *Uroviricota* whose putative hosts were categorized as Cyanobacteria (*i.e.,* cyanophages) along with a few whose hosts were *Alpha-* or *Beta-Proteobacteria* (Figure 5). In the control bottles, viral expression was dominated by a group of cyanophages distinctly separate from those in T_0_, indicating a moderate bottle effect on viral expression. The control bottles also saw a sharp expression increase in a group of 13 viral contigs, eight of which could be identified as RNA viruses classified in the phyla of *Duplornaviricota*, *Kitrinoviricota*, or *Negarnaviricota*. There was a wholesale shift in viral expression in the mitomycin C treatment. Here, viral expression was dominated by a group of 72 viral contigs, 40 of which could be categorized. Of these, 31 were identified as RNA viruses (mostly in the phylum *Kitrinoviricota*) and nine in the phylum *Uroviricota* with putative hosts likely *Alpha-* or *Beta-Proteobacteria*.

In the P-replete microcosms, differences between treatments were driven by virus groups and patterns distinct from those of the P-limited experiments. Expression of cyanophages was generally lower in the P-replete experiments (Figure 6). Viral expression in the T_0_ bottles was dominated by a group of 25 contigs, 14 of which could be categorized as *Uroviricota* with putative hosts distributed between *Cyanobacteria* and *Alpha-*, *Beta-*, and *Gamma-Proteobacteria*. Dominant viral expression in the control included the same contigs observed as dominant in T_0_, but with an additional group of 23 contigs most of which were phages whose putative hosts were *Gamma-Proteobacteria*. Again, there was a strong shift in expression observed in the mitomycin C treatment. Expression in mitomycin C bottles was dominated by 40 contigs, 21 of which could be categorized. Of these, eight were RNA viruses of the phyla of *Duplornaviricota*, *Kitrinoviricota*, or *Negarnaviricota*, while 13 were *Uroviricota* whose putative hosts were likely *Alpha-* or *Beta-Proteobacteria*. Details of viral classification is found in Supplemental Table 2.

### Mitomycin C is lethal to lab cultures of *M. aeruginosa*

We tested dose-dependent effects of mitomycin C on *M. aeruginosa* strains NIES-88 and NIES-298H. For both, the concentration of mitomycin C most commonly used in induction experiments (1 mg L^-1^) was lethal (Supplemental Figures 3A and 4). Concentrations of 0.1 mg L^-1^ inhibited population growth, but was not bactericidal. Mitomycin C at 0.01 mg L^-1^ weakly inhibited population growth of NIES-298H compared to control (Supplemental Figure 3A), but did not inhibit growth of NIES-88 (Supplemental 4). We tested two additional concentrations of mitomycin C in NIES-88. Mitomycin C at 0.5 mg L^-1^ was lethal to the population, while 0.05 mg L^-1^ was moderately inhibitory to growth (Supplemental Figure 3B).

In the filamentous cyanobacterium *Raphidiopsis* (*Cylindrospermopsis*) *raciborskii* Cr2010, 1 mg L^-1^ and 0.1 mg L^-1^ mitomycin C had strong inhibitory effects; neither were bactericidal (Supplemental Figure 5). A concentration of 0.01 mg L^-1^ had no inhibitory effect. In the filamentous cyanobacterium *Planktothrix agardhii* SB1031, 1 mg L^-1^ mitomycin C was strongly inhibitory but was not bactericidal (Supplemental Figure 6). Concentrations of 0.1 mg L^-1^ and 0.01 mg L^-1^ had little inhibitory effect (Supplemental Figure 6).

## Discussion

### The effects of mitomycin C

Mitomycin C is an antibiotic that cross-links complementary strands of DNA by covalently linking guanine nucleosides at CpG sites (46, 47). This cross-linking inhibits DNA replication and transcription at the site (48). A single cross-link per genome within an essential gene can be lethal in a bacterial cell (46). Cross-linking initiates DNA repair responses which can trigger prophage induction. Sensitivity of bacteria to mitomycin C varies by strain and species and can depend on specific traits such as GC content (48), presence of efflux pumps (49), and cellular oxidation status (46). The effects of mitomycin C range from bacteriostatic to bactericidal in a dose-dependent way. The concentration differences leading to bacteriostatic versus bactericidal effect can vary 2-to 7-fold in some species, while other species show no concentration difference (48).

It follows that if sensitivity to mitomycin C is dose-dependent, then induction of prophage is likely dose-dependent as well (50). An overdose of mitomycin C can be inhibitory and/or bactericidal before production of progeny virions is complete, while an underdose might fail to induce prophage. Both conditions can lead to an underestimation of lysogeny (43).

Within a community, application of a given concentration of mitomycin C will be an appropriate prophage-inducing dose to some and either an overdose or underdose to others, with researchers having little to no foreknowledge of the differential and selective effects of mitomycin C on the various components of the community. Here, we used metatranscriptomics to monitor the differential effects across community members. This method allowed us to detect and demonstrate that the standard protocol of 1 mg L^-1^ of mitomycin C is a highly-lethal overdose to freshwater cyanobacteria and strongly selects for *Proteobacteria* in the community. This overdose concept weighs heavily in our interpretation of the induction results.

From our data, we conclude that 1 mg L^-1^ mitomycin C was too bactericidal to the *Microcystis*-dominated cyanobacterial community to allow the induction of cyano-prophage which may have been present and/or the formation of new virions. It is conceivable that mitomycin C induced a heavily lysogenized community that led to extensive lysis, or to a cascade of lytic infections by progeny virions, but the results of direct viral counts seem to eliminate this as a possibility.

There are no data in the literature on mitomycin C dose effects on *Microcystis*. The standard and most commonly used concentration in both marine and freshwater induction studies is 1 mg L^-1^ (42). In the two previous induction studies conducted in *Microcystis* blooms, one used 1 mg L^-1^ (45), while the other used 20 mg L^-1^ (44). Consistent with our findings, neither study detected lysogeny in the *Microcystis*-dominated communities. Sulcius et al. (44) used the high concentration of 20 mg L^-1^ following observations of Dillon and Parry (51) in freshwater *Synechococcus* spp.

Dillon and Parry (51) tested 19 non-axenic strains of phycocyanin-rich freshwater *Synechococcus* for lysogeny using mitomycin C concentrations ranging from 1 to 100 mg L^-1^. They incubated cultures with mitomycin C for up to 14 d. They found that 16 strains were inducible and that 20 mg L^-1^ yielded the highest number of inductions. This high concentration induced a number of strains that were not inducible using 1 or 2 mg L^-1^. As controls, non-lysogenic strains were incubated with 20 mg L^-1^ mitomycin C with no visible lysis of cells at 14 d. Their results demonstrated high resistance to mitomycin C in some freshwater strains of *Synechococcus* spp.

Our results demonstrated the opposite in *Microcystis*. Metatranscriptomic analysis indicated that 1 mg L^-1^ mitomycin C incubated for 48 h effectively shut down transcription in natural populations of *Microcystis*. Follow-up lab studies testing effects of mitomycin C demonstrated that in cultures of *M. aeruginosa*, growth can be inhibited by concentrations as low as 0.01 and 0.05 mg L^-1^.

Transcriptional activity hinted that *Microcystis* was more sensitive to mitomycin C than other genera of commonly encountered freshwater cyanobacteria. Our lab studies were too limited to allow us to draw broad conclusions, but the results were consistent with metatranscriptomic observations from the field. Both *Raphidiopsis* and *Planktothrix* demonstrated greater resistance to mitomycin C than did *M. aeruginosa*, as measured by growth inhibition. This provides some measure of validation on our metatranscriptomic approach.

Our study reinforces, using specific freshwater taxa, what is already more generally known: the effects of mitomycin C are dose-dependent and vary between species (48). It seems safe to assume induction of freshwater lysogens is dose-dependent and varies between species as well. Furthermore, our observations indicate that the “correct” dosage (43) for induction-based lysogeny studies in natural *Microcystis* communities may be as low as 0.05 to 0.1 mg L^-1^.

### Lysogeny in Microcystis blooms

Longitudinal studies of cyanophage gene expression (and thus infection activity) suggest that lysogeny may be transiently prevalent in *Microcystis* over the course of a bloom (31, 41). Lysogeny offers an intriguing potential explanation of how *Microcystis*-dominated blooms sometimes seem at odds with ecological principles that appear to apply in other aquatic communities, e.g., “Paradox of the Plankton” (33) and “Kill-the-Winner” (36).

Based on a collection of early induction studies in marine systems, lysogenic infection was thought to predominate under conditions of low host abundance, low primary productivity, oligotrophic conditions, or in otherwise generally unfavorable conditions, while lytic infections predominated in near opposite conditions (9, 52). Under this model, *Microcystis* blooms would seem an unlikely environment in which to find a prevalence of lysogenic infections. A more recent study suggested that lysogenic infections can also predominate under high host densities (53), while a meta-analysis of 39 induction studies found no significant relationship between fraction of chemically inducible cells (FCIC) and host density (43). Knowles et al. (43) ultimately concluded that the constrained distribution of FCIC suggests that an as of yet unexamined variable, possibly environmental, may control dynamic prevalence of lysogenic infection. This would seem to reopen the door to lysogeny as an attractive hypothesis to help explain dynamics of *Microcystis* blooms. Existence of transient lysogeny could provide homoimmunity, which in turn could play a pivotal role in the ecology and dynamics of freshwater harmful algal blooms. To our knowledge, direct evidence demonstrating widespread lysogeny in natural *Microcystis* populations is lacking. But as the maxim says, absence of evidence is not evidence of absence, and the gravity of the topic argues for continued research in this area. The challenge, then, is to design and conduct experiments that can quantitatively detect lysogeny in natural blooms. Here, using community metatranscriptomics, we demonstrate that the standard protocol for detecting lysogeny in natural aquatic communities is inappropriate for *Microcystis* research.

## Conclusion

Evidence indicates that viruses are important in the ecology of *Microcystis*, while cell concentrations during blooms seemingly contradict tenets of ecological models of viral infection in phytoplankton. Lysogeny in *Microcystis* could help explain this phenomenon. Consistent with this idea, metatranscriptomic studies suggest that lysogeny is dynamic during blooms. The challenge in testing this hypothesis is to quantitatively detect lysogeny in bloom communities. Application of mitomycin C is a time-honored technique used to detect lysogeny in phytoplankton. We used mitomycin C to test for lysogeny in a *Microcystis* bloom and with metatranscriptomic analysis demonstrated that the standard protocol of 1 mg L^-1^ was a highly-toxic overdose which likely inhibited induction of any prophage present. Follow-up lab studies indicate that 0.1 mg L^-1^ may be more appropriate in *Microcystis*. These findings will guide future efforts to detect lysogeny in blooms, which in turn is needed to understand the role of lysogeny in the ecology of *Microcystis*. The detrimental effects of HABs on freshwater ecosystems argues for such a gain in understanding.

## Methods

### Lake Erie microcosm experiments

The mitomycin C field experiments reported here were a subset of a larger microcosm study conducted at Ohio State University’s Stone Laboratory in Put-in-Bay, Ohio in 2019. Detailed descriptions of the larger studies were reported in Pound et al. (54) and Martin et al. (55).

Microcosm experiments were independently conducted twice with natural communities collected on different days and from different locations in Lake Erie. Surface water was collected for the first experiment in 20-L carboys at 41° 44.946 N, 83° 06.448 W in Lake Erie on 21 July 2019. Water for the second experiment was collected at 41° 49.568 N, 83° 11.678 W on 24 July 2019. Sampling at both sites occurred during an early-phase *Microcystis*-dominated bloom (56). Physicochemical measurements of surface water conditions were made with a YSI EXO2 sonde at the time of water collection.

For each of the mitomycin C experiments, microcosms were established by aliquoting homogenized lake water containing natural communities into nine 1.2-L polycarbonate bottles, which were randomly allocated between three treatments with three replicate bottles per treatment: mitomycin C (1 mg L^-1^), initial conditions (T_0_), and control. The three T_0_ bottles were sampled immediately. The control and mitomycin C bottles were incubated *in situ* for 48 h in Lake Erie in floating corrals off the dock at the Stone Lab (41° 39.467 N, 82° 49.600 W). The corrals were covered with shade screen which reduced incident photosynthetically active radiation by ∼40%.

Bottles were sampled for chl *a* concentration, RNA sequencing, and direct viral counts. For chl *a* samples, 100 mL of water was filtered through 0.2-µm-pore-size polycarbonate filters. chl *a* was extracted from the filters in 90% acetone at 4° C for 24 h then quantified on a 10-AU Fluorometer (Turner Designs) following the method of N. A. Welschmeyer (57). RNA samples were collected by filtering ∼150 mL of water through 0.2-µm-pore-size Sterivex^TM^ filters then flash freezing in liquid nitrogen. Samples were stored at -80° C until extraction. For virus count samples, 5 mL of lake water was flash frozen in liquid nitrogen and stored at -80° C until enumeration.

Chl *a* response indicated that the phytoplankton community in the first experiment was P-limited while the community in the second experiment was P-replete (see Fig. 2 in Martin et al. (55)). This fortuitously allowed us to compare the effects of mitomycin C between P-limited and P-replete communities. Results from the two experiments are presented separately and are referred to as P-limited and P-replete.

### Direct viral counts

VLPs were enumerated following the procedures summarized in E. R. Gann et al. (58) and C. P. Brussaard (59). Raw lake water was prefiltered using 0.45-µm pore-size polyvinylidene difluoride syringe filters (Millipore Sigma). Prefiltered lake water was then fixed with 0.5% glutaraldehyde at 4° C for 30 min. Fixed viruses were stained with 1x concentration of SYBR Green I DNA stain (Lonza Bioscience) and incubated at 80° C for 10 min. Stained viruses were enumerated on a FACSCalibur (BD Biosciences) flow cytometer gating on SYBR green emission (520 nm) and side scatter. Known concentrations of 1-µm yellow-green FluoSphere Carboxylate-Modified Microspheres (505/515 nm) (Invitrogen) were added to samples to provide for absolute quantification of VLPs.

### RNA extraction and sequencing

RNA was extracted from Sterivex filters as described on *protocols.io* (60). Briefly, RNA was extracted with acid phenol and chloroform, precipitated with sodium acetate and 100% ethanol, and washed with 70% ethanol. Residual DNA was removed by digestion using a Turbo DNA-*free* Kit (Ambion) following the protocol on *protocols.io* (61). RNA samples were considered DNA free if no bands were visible on an agarose gel after 30 cycles of PCR amplification using standard primers 27F/1522R targeting the 16S rRNA gene (62). Samples showing DNA contamination were retreated with Turbo DNA until no bands were visible. RNA was quantified using the Qubit hsRNA assay.

cDNA libraries were prepared at Discovery Life Sciences (Huntsville, Alabama) using the Illumina Stranded Total RNA Prep, Ligation with Ribo-Zero Plus kit. Libraries were sequenced at Discovery Life Sciences on the Illumina NovaSeq platform generating ∼100-120 million 100-bp PE reads per library.

### Bioinformatic analysis

Residual ribosomal reads were removed *in silico* using BBDuk (v. 38.90) in the BBTools package with the Silva database (v. 119) as the ribosomal sequence reference (63, 64). Reads were trimmed for quality using CLC Genomics Workbench (v. 20.0.4) using a quality limit score of 0.02, ambiguous nucleotides = 0, and minimum length = 50 bp. All other settings were default values. Nonribosomal, trimmed reads from all libraries within an experiment were combined and assembled together into a single co-assembly using MegaHit (v. 1.2.9) (65), *i.e.*, a separate co-assembly was produced for the P-limited and P-replete experiments.

For viral community analysis, viral contigs in the co-assemblies were identified using VirSorter2 (v. 2.2.3) (66). Contigs identified as viral by VirSorter2 were further analyzed with CheckV (v. 0.8.1) for additional verification of viral origin and for viral genome completeness (67). Contigs labeled as “no virus genes detected” in CheckV were removed from the putative viral contig list and all downstream analysis. Remaining viral contigs were classified taxonomically using the Contig Annotation Tool (CAT) (68). DIAMOND (v. 2.0.14) was used to identify best protein alignment hits to amino acid translations of genes predicted in CAT (*via* Prodigal v. 2.6.3) (69, 70). Categorization of putative hosts of the *Uroviricota* (the tailed bacteriophages) was made by a manual decision when the best DIAMOND hit (with a E-value cutoff of 1 x 10^-99^) of more than one ORF in a contig was to a characterized phage with a known host. This approach allowed us to place phage contigs into a conservative classification system of being either cyanophage or phage likely infecting heterotrophic bacteria. Contigs with hits higher than the cutoff or to phages with ambiguous or unknown hosts were left un-categorized.

Quantification of DNA virus infection activity and presence/activity of RNA viruses in each treatment was estimated by mapping reads from each library to viral contigs. Reads were mapped in CLC Genomics Workbench using settings of 0.85 and 0.85 for length fraction and similarity fraction, respectively. Default settings were used for other parameters. Expression was calculated as transcripts per million (TPM) (71) with reads mapped as pairs counted as two and reads mapped as broken pairs counted as one. Expression of each putative viral contig was standardized across replicates and treatments. Heatmaps illustrating standardized viral expression/presence across treatments were made with Heatmapper (72) clustering contigs using average linkage with Pearson distance. By sequencing RNA, we captured infection activity of DNA viruses and presence and/or infection activity of RNA viruses. For economy’s sake, we will refer collectively to presence/activity of either virus type as “viral expression” or “viral activity”.

To estimate community gene expression, putative genes within a co-assembly were first identified using MetaGeneMark (v. 3.38) (73). Identified genes were annotated with predicted function and taxonomy using GhostKOALA (74). The resulting final gene list used in downstream analysis included only those genes with predicted taxonomy. Expression in TPM for each gene was calculated by mapping reads from each library to the gene list using the same parameters described for viral expression.

### Laboratory experiments, strains, and culturing

Axenic cultures of *Microcystis aeruginosa* strains NIES-88 and NIES-298 were grown in 50-mL glass tubes in CT medium (75) modified by supplying P at an equivalent molar concentration via Na_2_HPO_4_ rather than the original Na_2_-B-glycerophosphate. NIES-88 was purchased from the Microbial Culture Collection of the National Institute for Environmental Studies (Japan). NIES-298 was provided by Jozef Nissimov (University of Waterloo). These *Microcystis* strains were used because of their relevance in mitomycin C sensitivity tests. The NIES-88 genome harbors a contiguous viral segment ∼37 kbp long (31), which is assumed to be a remnant of a defective prophage, while NIES-298 is the host of the cyanophage Ma-LMM01 (20). Non-axenic cultures of *Planktothrix agardhii* SB-1031 and *Raphidiopsis* (*Cylindrospermopsis*) *raciborskii* Cr2010 were grown in 50-mL glass tubes in standard MLA medium (76). SB1031 was isolated from Sandusky Bay, Lake Erie and provided by George S. Bullerjahn (Bowling Green State University). Cr2010 was isolated from Reeuwijkse Plassen in the Netherlands (77) and was provided by Corina Brussaard (Royal Netherlands Institute for Sea Research). All cultures were grown at 26° C with a photosynthetic fluence rate of ∼50 µmol photons m^-2^ s^-1^ provided by cool-white fluorescent bulbs (GE Ecolux 32W) on a 12-h light/dark cycle. Temperature was measured every 30 min using a Hobo Tidbit TempLogger (OnSet Computer Corporation).

To test sensitivity to mitomycin C, cultures were grown across a series of mitomycin C concentrations. Mitomycin C (Thermo Fisher Scientific) was dissolved in DMSO then added to cultures to produce final test concentrations of 0.01, 0.1, and 1 mg L^-1^. A DMSO solvent-only control and a no solvent/no mitomycin C control were included. In experiments using *M. aeruginosa* NIES-88, we tested the additional mitomycin C concentrations of 0.05 and 0.5 mg L^-1^. Chl *a* fluorescence was used as a proxy for cyanobacterial biomass and was measured on a TD-700 fluorometer (Turner Designs). All experiments were conducted in biological triplicate.

## Data availability

Raw reads are publicly available in the Sequence Read Archive of the National Center for Biotechnology Information under the BioProject numbers PRJNA737197 and PRJNA823389.

## Acknowledgments

This work was supported by funding from the *Simons Foundation* (735077) as well as NIEHS (1P01ES028939–01) and NSF (OCE-1840715) through the *Great Lakes Center for Fresh Waters and Human Health* at *Bowling Green State University.* We also acknowledge the *Kenneth & Blaire Mossman Endowment* at the *University of Tennessee*.

## Author Contributions

SWW conceptualized and designed the experiments. HLP, RMM, and JDC conducted the microcosm experiments. HLP extracted/processed samples. EAB conducted growth curves. RMM and ERD analyzed data. RMM and SWW wrote the original draft of the manuscript. All authors participated in revision of and accepted final version of the manuscript.

## Supplemental Information

**Supplemental Figure 1.**
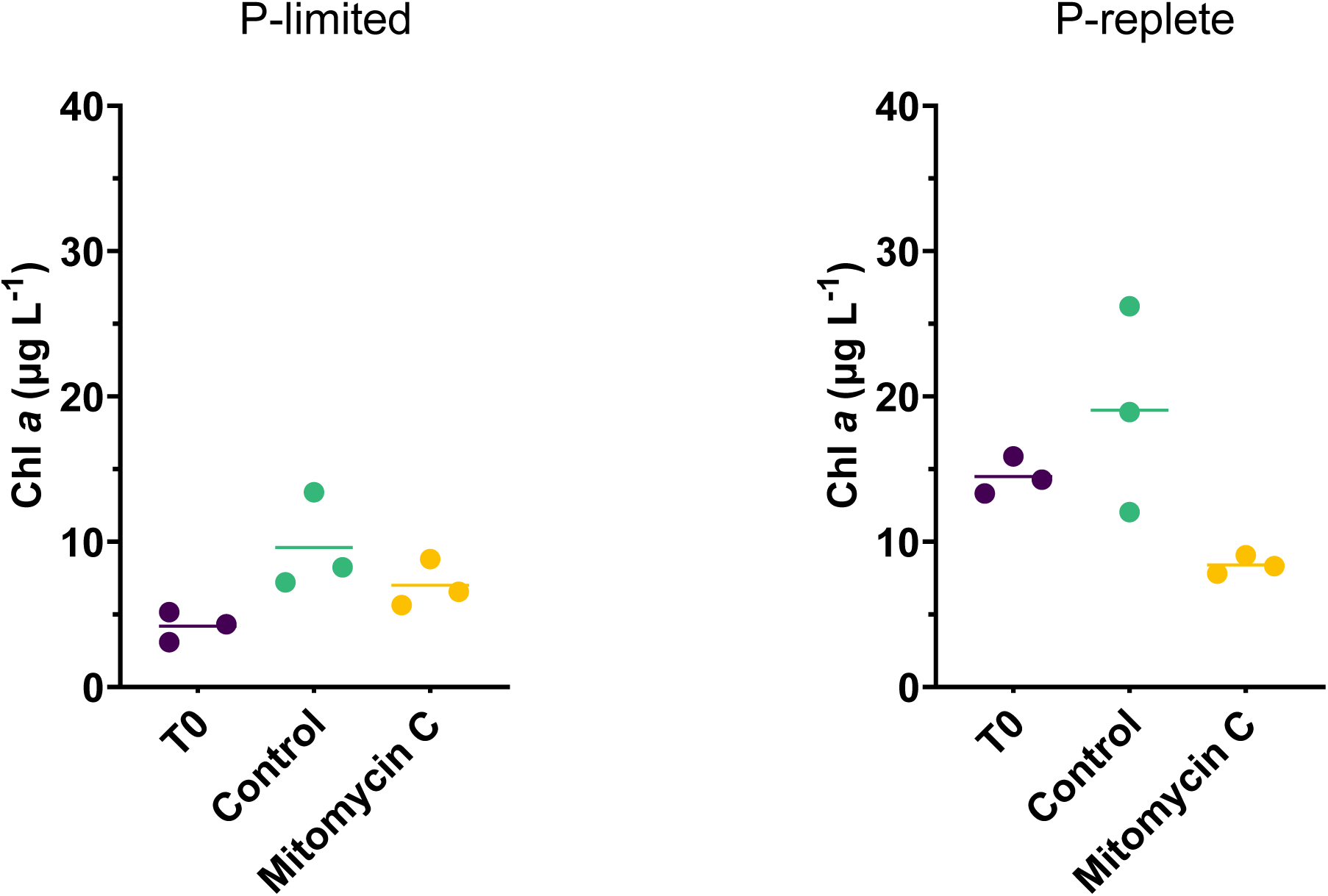
Chlorophyll *a* response by treatment. a) P-limited experiment. b) P-replete experiment.

**Supplemental Figure 2.**
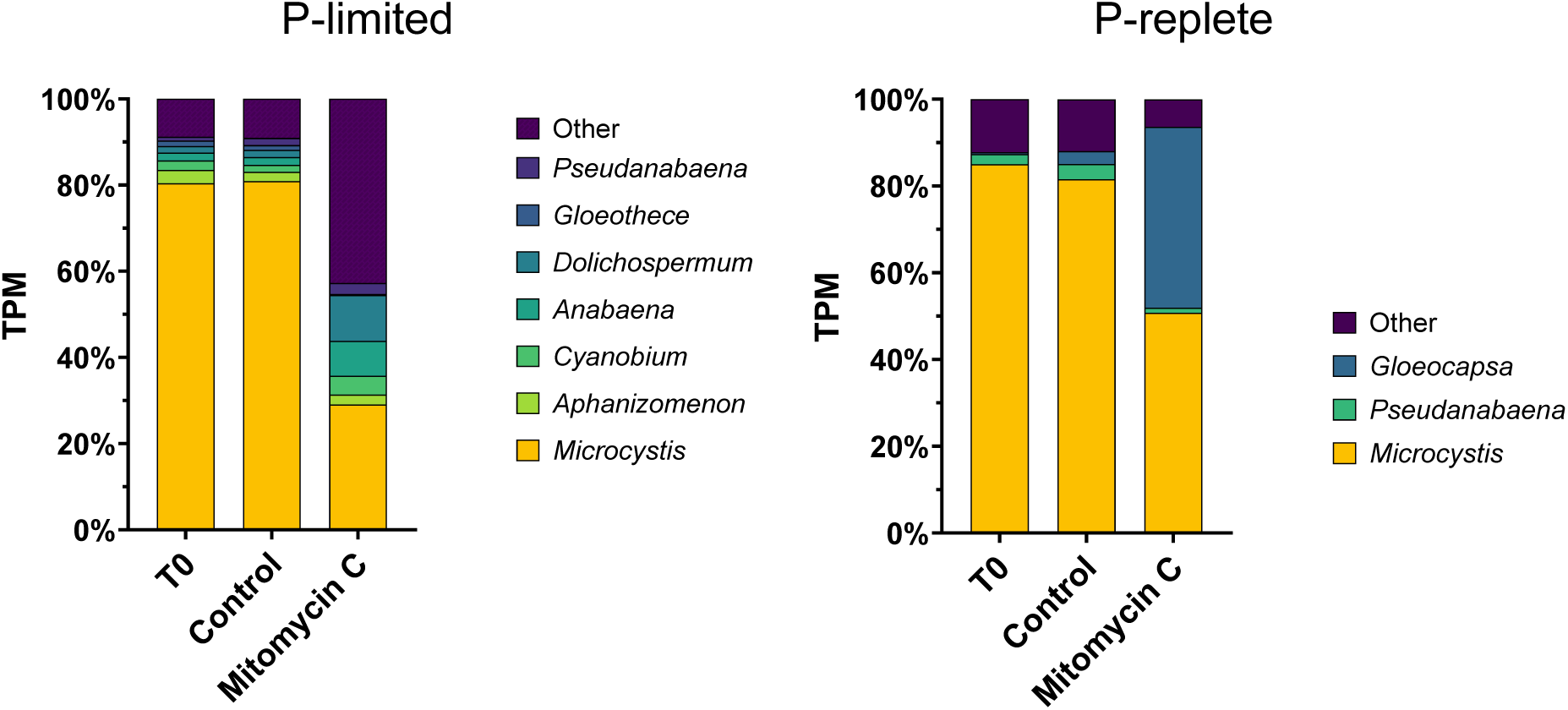
Transcription activity of major genera of Cyanobacteria Phylum by treatment as a percent of total cyanobacterial transcription. a) P-limited experiment. b) P-replete experiment.

**Supplemental Figure 3A.**
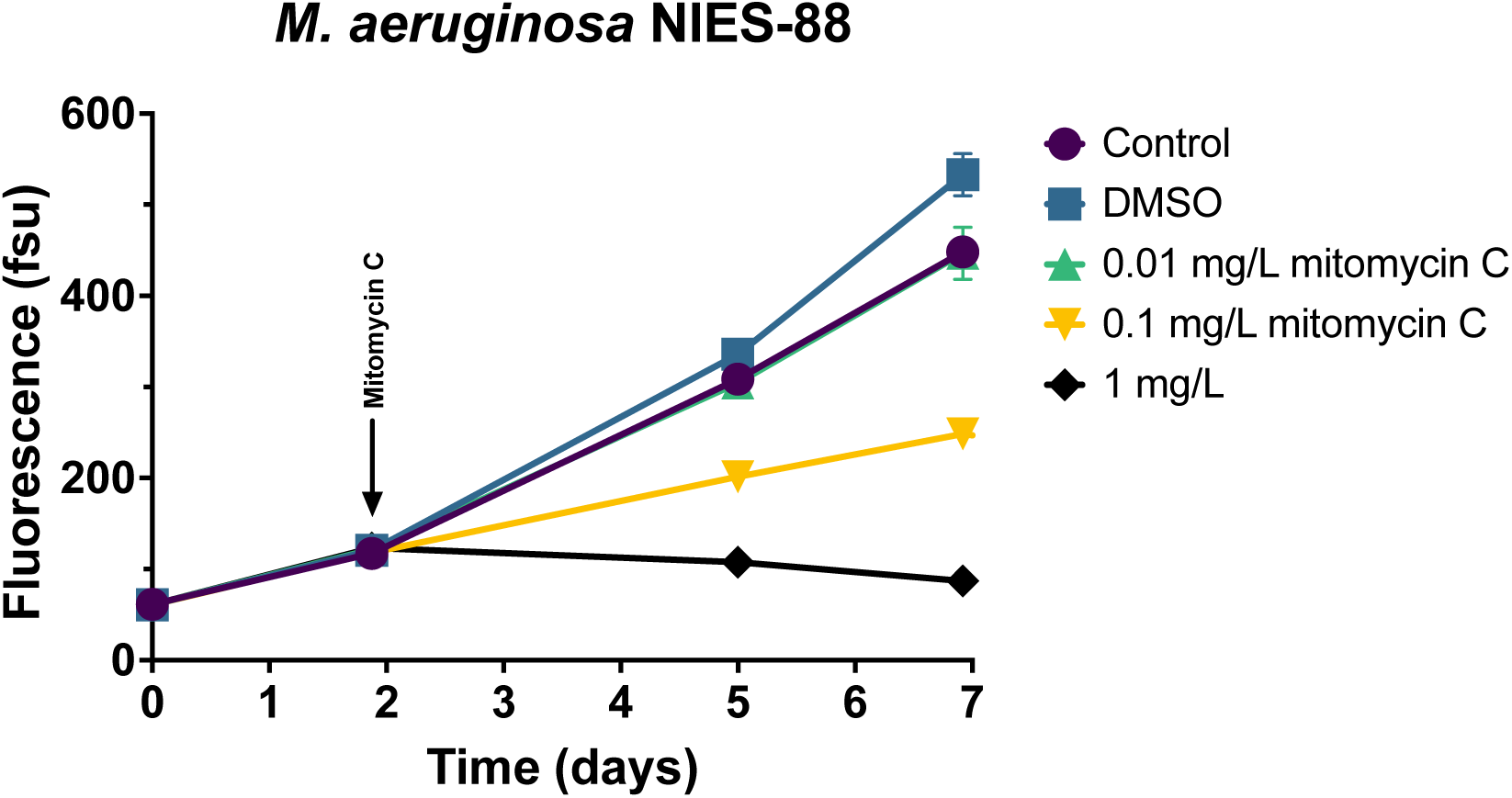
Dose-dependent growth response of *Microcystis aeruginosa* NIES-88 to mitomycin C.

**Supplemental Figure 3B.**
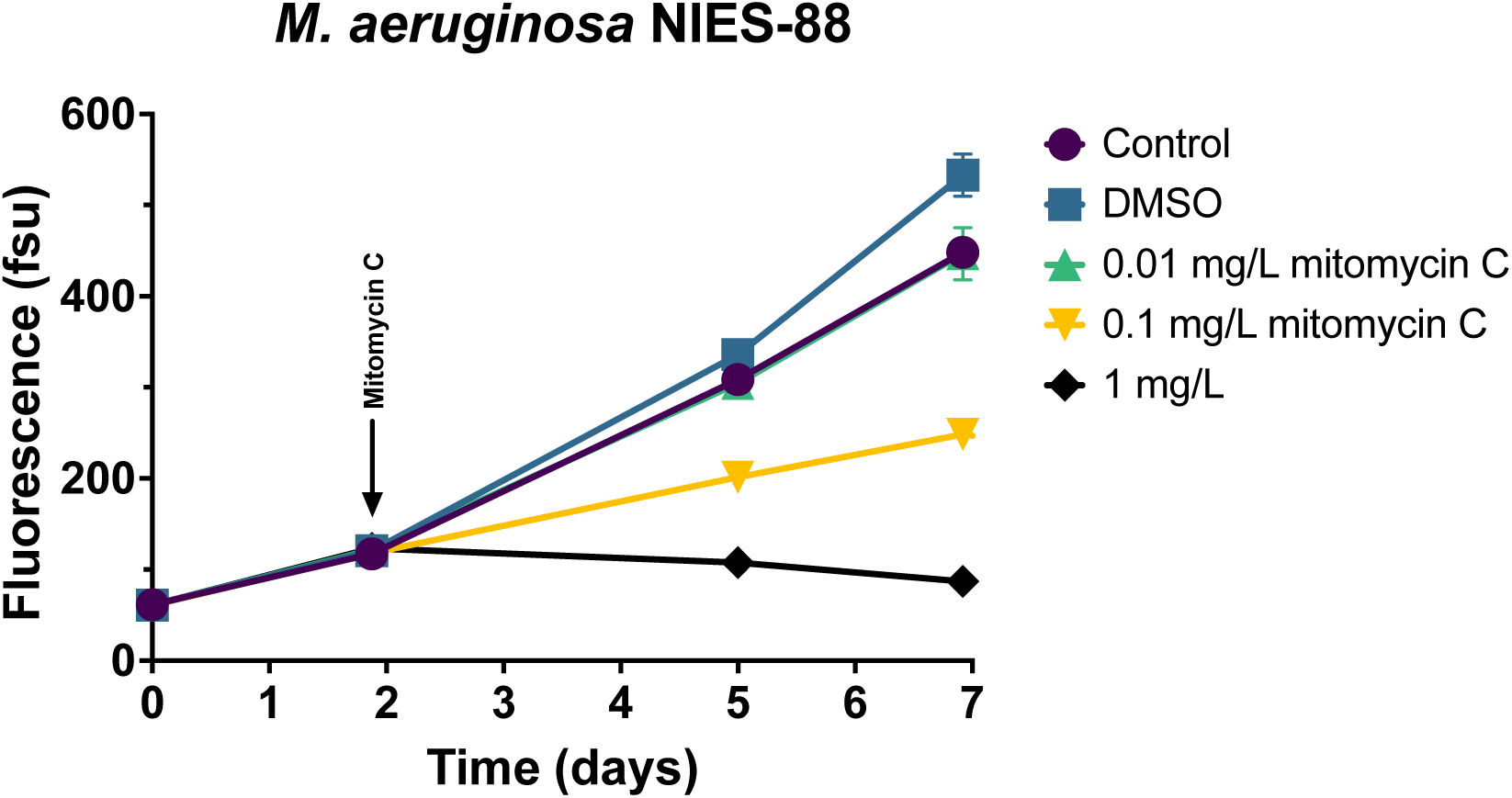
Dose-dependent growth response of *Microcystis aeruginosa* NIES-88 to mitomycin C.

**Supplemental Figure 4.**
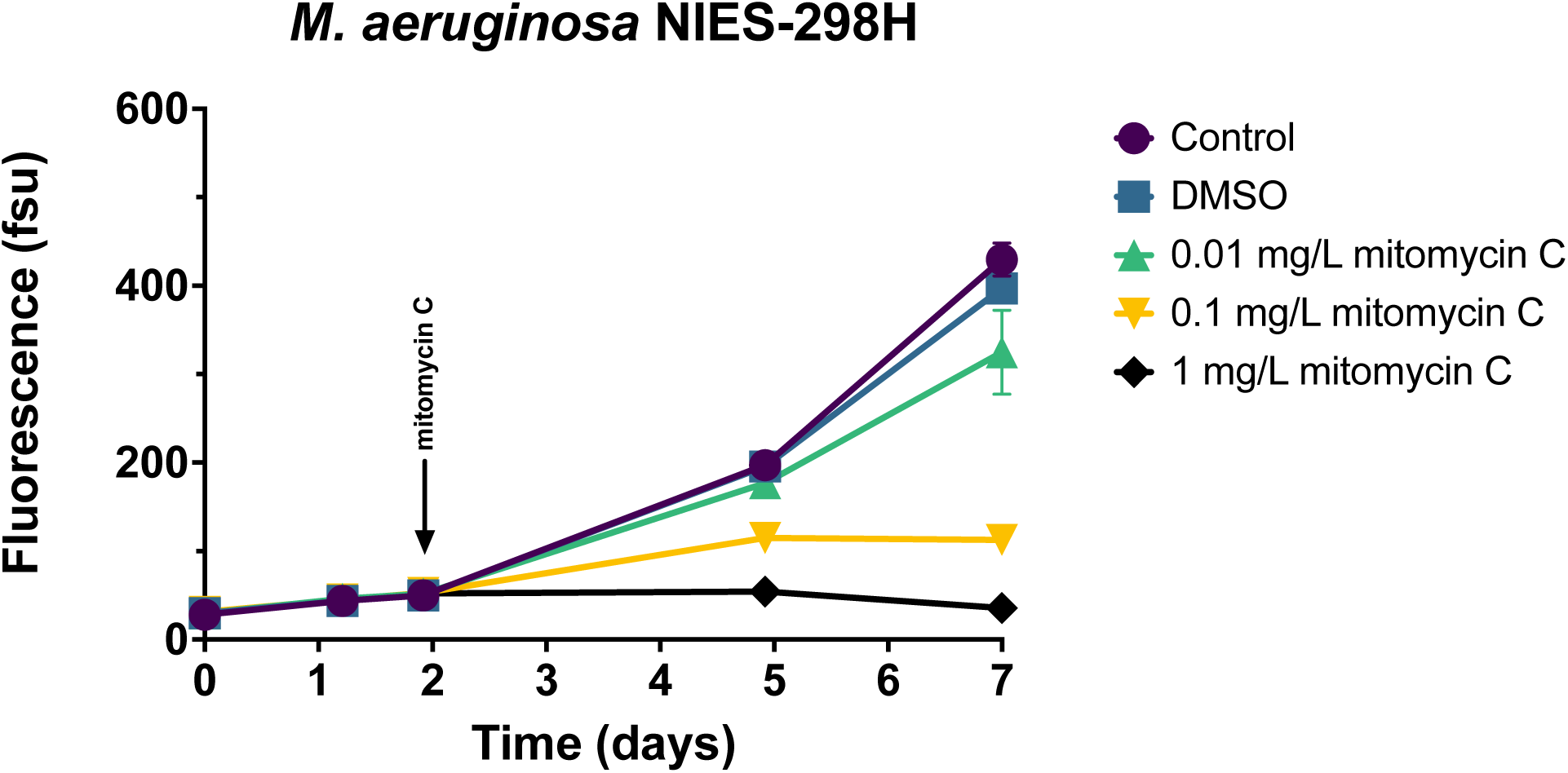
Dose-dependent growth response of *Microcystis aeruginosa* NIES-298H to mitomycin C.

**Supplemental Figure 5.**
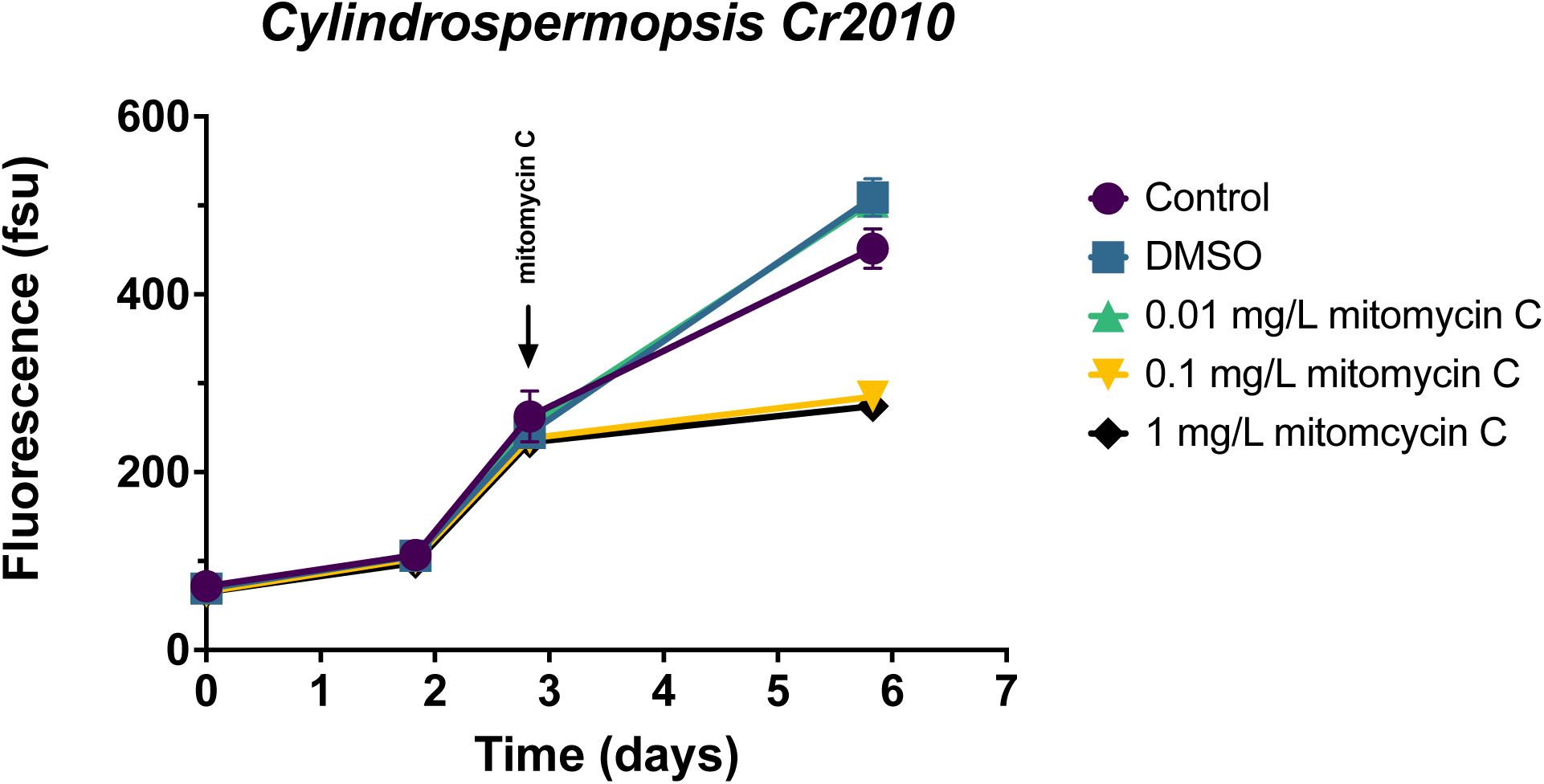
Dose-dependent growth response of *Raphidiopsis (Cylindrospermopsis) raciborskii* Cr2010 to mitomycin C.

**Supplemental Figure 6.**
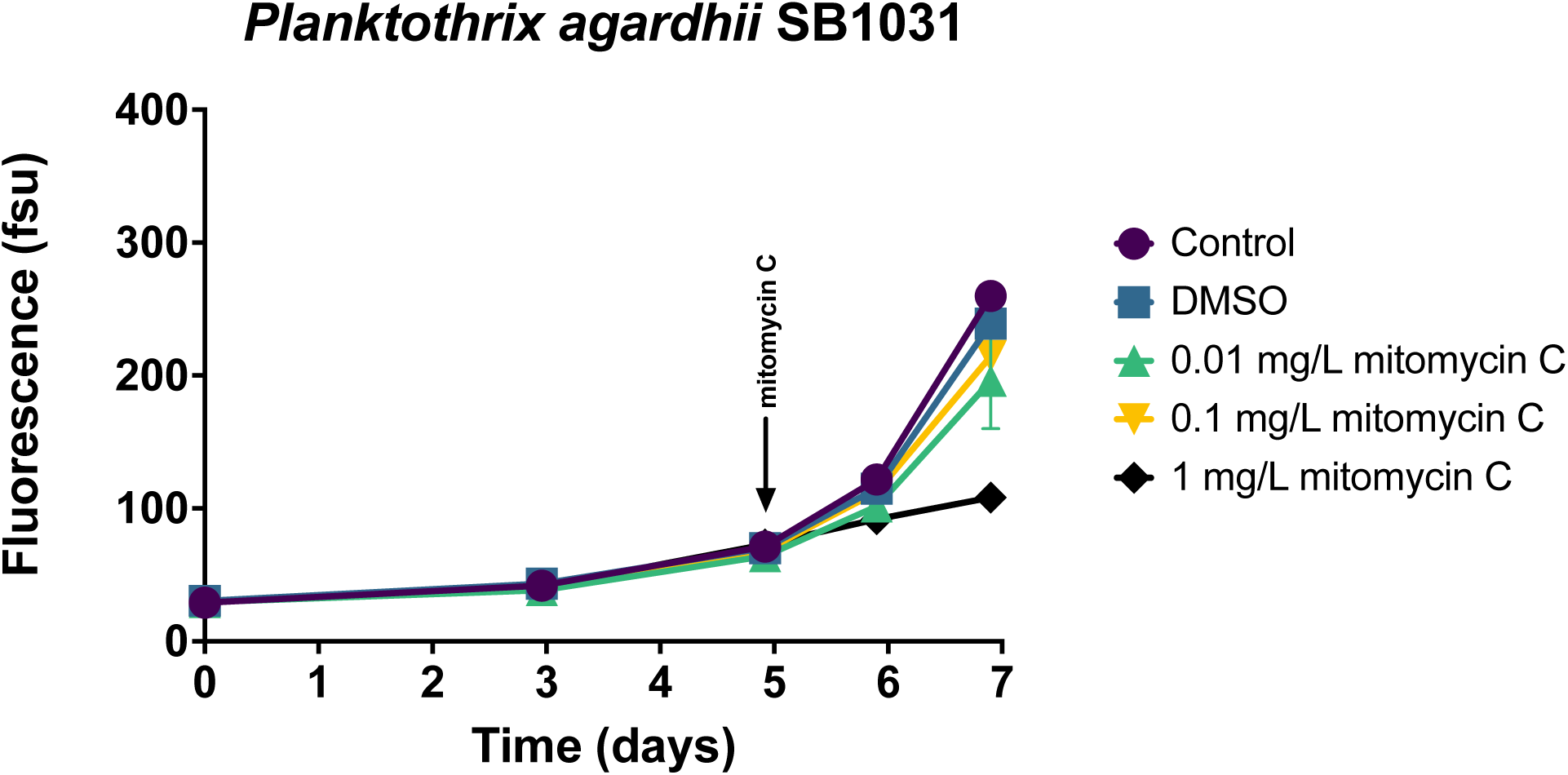
Dose-dependent growth response of *Planktothrix agardhii* SB1031 to mitomycin C.

Supplemental Table 1. Attached as Excel spreadsheet.

Supplemental Table 2. Attached as Excel spreadsheet.

